# Highly multiplexed immune profiling throughout adulthood reveals kinetics of lymphocyte infiltration in the aging mouse prostate

**DOI:** 10.1101/2020.06.18.160556

**Authors:** Jonathan J. Fox, Takao Hashimoto, Héctor I. Navarro, Alejandro J. Garcia, Benjamin L. Shou, Andrew S. Goldstein

## Abstract

Aging is a significant risk factor for cancer in several tissues, including the prostate. Defining the kinetics of age-related changes in these tissues is critical for identifying regulators of aging and evaluating interventions to slow the aging process and reduce disease risk. An altered microenvironment is characteristic of prostatic aging in mice. Whether features of aging in the prostate emerge predominantly in old age or earlier in adulthood has not previously been established. Using comprehensive immune profiling and time-course analysis, we show that populations of T and B lymphocytes increase in the mouse prostate between 6 and 12 months of age. When comparing the prostate to other urogenital tissues, we found similar features of age-related inflammation in the mouse bladder. In summary, our study offers new insight into the kinetics of prostatic inflammaging and the window when interventions to slow down age-related changes may be most effective.

## INTRODUCTION

Aging is associated with the dysfunction of many different cellular and molecular processes which results in the development of age-related pathologies including cancer (Lopez-Otin et al., 2013). The risk of prostate cancer increases with age (Leitzmann and Rohrmann, 2012), but the mechanisms driving increased disease risk with age are poorly understood. One well-established aspect of aging is the development of chronic low-grade inflammation, termed “inflammaging” (Franceschi et al., 2018). Chronic inflammation in the benign prostate increases the risk for developing high-grade prostate cancer later in life (Gurel et al., 2014). Prostatic inflammation is associated with luminal progenitor cells that can serve as target cells for human prostate cancer initiation (Liu et al., 2016). In both mice and humans, the population of luminal progenitor cells in the prostate increases with age (Crowell et al., 2019), expanding the pool of potential target cells for transformation.

While mice do not naturally develop prostate cancer, alterations in the hormonal milieu and genetic alterations in the epithelium can cooperate with aging to promote tumorigenesis (Grabowska et al., 2014). Importantly, the cellular changes that define aging phenotypes in the human prostate are conserved in the aging mouse prostate, indicating that the mouse is an ideal model system to investigate factors that regulate age-related changes to the prostate (Crowell et al., 2019). The aging mouse prostate is characterized by increased inflammation and stromal disorganization (Bianchi-Frias et al., 2010). Compared with sexually mature 3-month-old mice, prostates of 24-month-old mice are enriched for T and B lymphocytes. Whether prostatic inflammation emerges primarily in old age or steadily increases throughout adulthood has not been established.

Efforts to slow age-related changes have garnered considerable interest over the past decade. Caloric restriction, as well as mTOR inhibition, show promise in delaying aging phenotypes and extending lifespan in model organisms (Heilbronn and Ravussin, 2003; Johnson et al., 2013). Pharmacological enhancement of NAD^+^ levels has also been shown to reverse aging phenotypes (Rajman et al., 2018). Anti-inflammatory strategies represent another option for combating the effects of aging (Pedersen, 2009).

Understanding the dynamics of age-related changes is essential for defining the window in which interventions are most efficacious.

We sought to define the kinetics of mouse prostatic inflammation throughout adulthood at single-cell resolution using mass cytometry, or cytometry by time-of-flight (CyTOF), which has been previously reported by our group and others to study immune cells in the prostate (Crowell et al., 2019; Fox et al., 2019; Wang et al., 2016). We also set out to determine how aging-associated inflammation in the prostate compares with other urogenital tissues susceptible to age-related cancers Our study revealed features of old age in the prostate far earlier than previously reported, with evidence of increased lymphocyte infiltration emerging between 6 and 12 months of age. Age-related changes in immune profiles were shared by the mouse prostate and bladder, suggesting common features of inflammaging in distinct genitourinary tissues.

## RESULTS

### Identification of distinct immune subsets in mouse prostate using mass cytometry

Upon evaluating prostates from mice at various ages between 1 and 16 months, we observed a positive correlation between age and wet prostate weight (ρ = 0.6673, P_ρ_ < 0.001) characterized by a rapid increase in weight in pre-pubescent mice (< 3 months old) (Figure S1A). This growth model is consistent with the rapid, androgen-dependent growth of the prostate which occurs postnatally until sexual maturity at 2–3 months of age (Dutta and Sengupta, 2016; Sugimura et al., 1986). Our data also demonstrates that the mouse prostate continues to grow after reaching adulthood. Analyzing post-pubertal adult mice from 3 to 16 months of age, we observed a positive correlation between age and prostate weight (ρ = 0.5144, P_ρ_ < 0.01), with a significant increase in mass at 16 months of age (p < 0.05) (Figure S1A). Age was also positively correlated with the percentage of CD45^+^ immune cells in the prostate, as measured by flow cytometry (ρ = 0.7034, P_ρ_ < 0.01) (Figure S1B), consistent with age-related prostatic inflammation.

We used CyTOF to characterize age-related changes in the immune microenvironment of the prostate throughout adulthood. Prostates were isolated from adult mice aged 3, 6, 9, 12, and 16 months (each n = 4), and dissociated cells were stained with a panel of 19 metal-tagged antibodies designed to label a broad variety of immune cell types (Table 1 and Figure 1A). Single live immune cells were concatenated and clustered as a whole to allow for comparisons of phenotypically-identical clusters using the t-distributed stochastic neighbor embedding (t-SNE) algorithm for dimensional reduction and visualization (Figures 1A and 1B and Figure S2A). Based on expression of immune cell lineage markers, the t-SNE plot was separated into 3 broad groups: T cells (expressing CD3e, CD4, or CD8a), B cells (expressing B220), and myeloid/NK cells (expressing CD335, F4/80, CD11b, CD11c, Ly6C, or Ly6G) (Figure 1B). Correlation analysis showed a positive correlation between age and the percentage of T cells (ρ = 0.8952, P_p_ < 0.0001) and B cells (ρ = 0.8216, P_ρ_ < 0.0001) in the adult mouse prostate (Figures 1C and 1D). The increase in T and B lymphocytes with age was accompanied by a respective negative correlation between age and the percentage of myeloid/NK cells (ρ = −0.9197, P_ρ_ < 0.0001) in the adult mouse prostate (Figure 1E). These trends are consistent with the age-related increase in T and B lymphocytes and decrease in myeloid and NK cells previously reported in 3- and 24-month-old mouse prostates by our group (Crowell et al., 2019).

**Figure 1.**
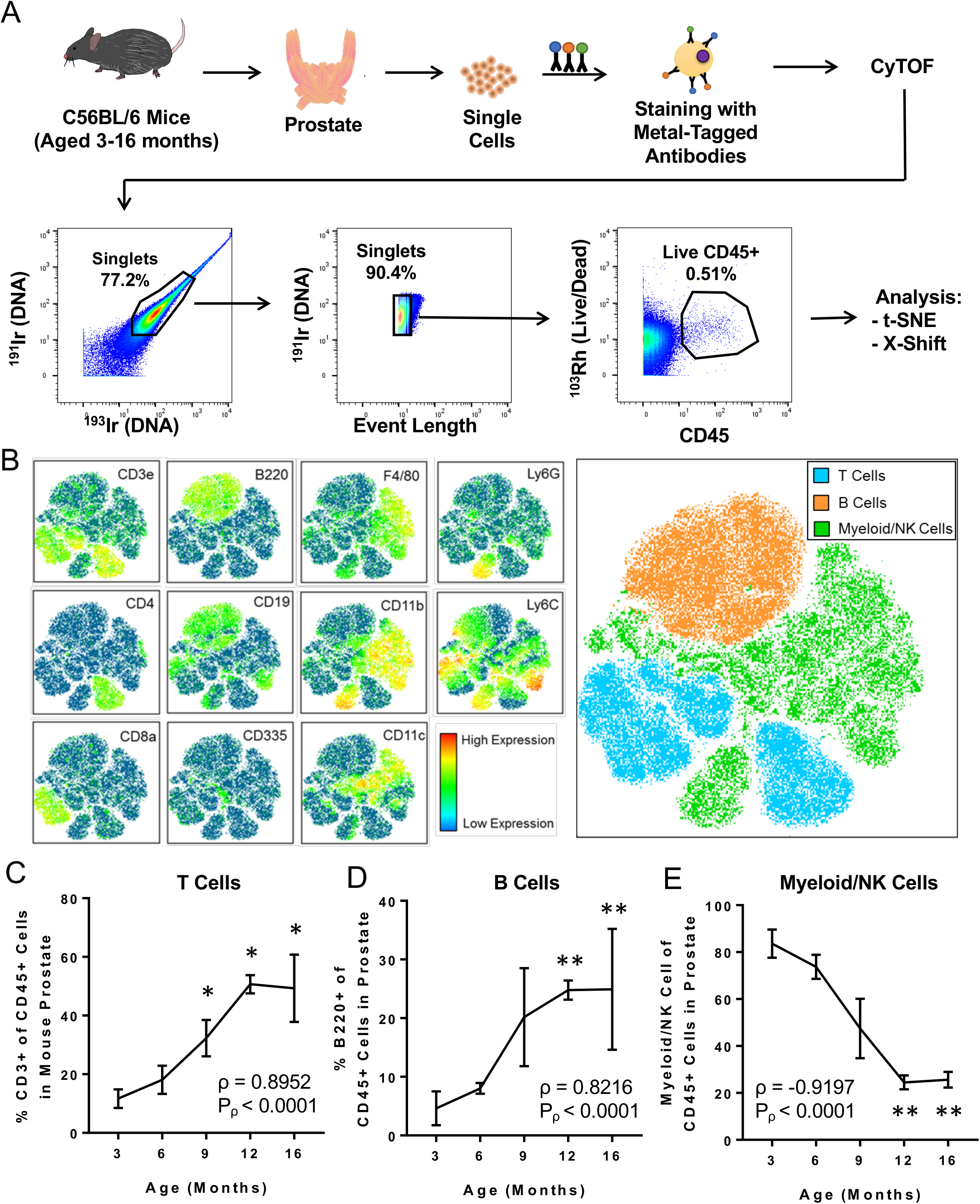
CyTOF immunophenotyping of the mouse prostate. (A) Workflow for prostate immunophenotyping using CyTOF. Mouse prostates of different ages were isolated, dissociated to single cells, and stained with a panel of heavy metal-tagged antibodies before data acquisition via mass cytometry. Bivariate plots show gating for single, live CD45+ immune cells before analysis using clustering algorithms (t-SNE and X-Shift). Percentages represent the fraction of events that are within the gate in each bivariate plot. ^191^Ir/^193^Ir labels DNA of all cells. ^103^Rh labels DNA of dead cells. This flowchart was adapted from a previous publication from our group (Crowell et al., 2019). (B) t-SNE plot generated from the immune cells from mouse prostate, bladder, and kidney. Left: Heat maps showing expression of selected lineage markers by immune cells clustered using t-SNE. Right: t-SNE plot separated into 3 broad groups of immune cells (T cells, B cells, and myeloid/NK cells). (C–E) Quantification of changes to the mouse prostate immune cell composition during adult aging for CD3^+^ T cells (C), B220^+^ B cells (D), and myeloid/NK cells (E). Spearman correlation coefficient (ρ) and associated p-value (P_ρ_) represent the correlation between % immune cell type and age. Data represents mean ± SD of 4 biological replicates at each age. Kruskal-Wallis, p < 0.01 (T cells, B cells, and Myeloid/NK cells). Dunn’s multiple comparisons test against 3-months-old, *p < 0.05, **p < 0.01. See also Figures S1 and S2.

**Table 1.** CyTOF immunophenotyping antibody panel for discovery experiment.

| Label | Target | Clone | Conjugation | Source |
| --- | --- | --- | --- | --- |
| <sup>89</sup>Y | CD45 | 30-F11 | Pre-conjugated | Fluidigm |
| <sup>139</sup>La | CD27 | LG.3A10 | Maxpar kit | BioLegend |
| <sup>141</sup>Pr | Ly6G | 1A8 | Pre-conjugated | Fluidigm |
| <sup>146</sup>Nd | F4/80 | BM8 | Pre-conjugated | Fluidigm |
| <sup>147</sup>Sm | CD80 (B7-1) | 16-10A1 | Maxpar kit | BioLegend |
| <sup>148</sup>Nd | CD11b | M1/70 | Pre-conjugated | Fluidigm |
| <sup>150</sup>Nd | Ly6C | HK1.4 | Pre-conjugated | Fluidigm |
| <sup>152</sup>Sm | CD3e | 145-2C11 | Pre-conjugated | Fluidigm |
| <sup>153</sup>Eu | CD274 (PD-L1) | 10F.9G2 | Pre-conjugated | Fluidigm |
| <sup>154</sup>Sm | CD152 (CTLA-4) | UC10-4B9 | Pre-conjugated | Fluidigm |
| <sup>155</sup>Gd | CD25 | 3C7 | Maxpar kit | BioLegend |
| <sup>156</sup>Gd | CD4 | RM4-5 | Maxpar kit | BioLegend |
| <sup>159</sup>Tb | CD279 (PD-1) | 29F.1A12 | Maxpar kit | BioLegend |
| <sup>166</sup>Er | CD19 | 6D5 | Pre-conjugated | Fluidigm |
| <sup>167</sup>Er | CD335 (NKp46) | 29A1.4 | Pre-conjugated | Fluidigm |
| <sup>168</sup>Er | CD8a | 53-6.7 | Pre-conjugated | Fluidigm |
| <sup>172</sup>Yb | CD86 (B7-2) | GL1 | Pre-conjugated | Fluidigm |
| <sup>176</sup>Yb | CD45R (B220) | RA3-6B2 | Pre-conjugated | Fluidigm |
| <sup>209</sup>Bi | CD11c | N418 | Pre-conjugated | Fluidigm |

To provide more granularity to our analysis we performed unsupervised k-nearest neighbors clustering using the X-Shift algorithm to further subdivide the t-SNE plot into phenotypically-distinct immune cell clusters (Figure 2A and Figure S2B). The X-Shift algorithm was chosen because it optimizes the numbers of clusters to prevent over- and under-clustering (Samusik et al., 2016). The algorithm produced 29 immune cell clusters which were then subdivided by cell type. Expression of lineage markers was used to identify 24 of 29 immune cell clusters as T cells (CD3e+), B cells (B220+ and CD19+), NK cells (CD335+), and myeloid cells (any combination of F4/80+, CD11b+, CD11c+, or Ly6G+) (Figures 2A–2E). The 5 remaining clusters were classified as unknown due to expression of both T cell and myeloid cell markers (U1), lack of a strong signal for any lineage-specific markers (U2–U3, U5) or mild staining for every marker (U4) (Figure 2A).

**Figure 2.**
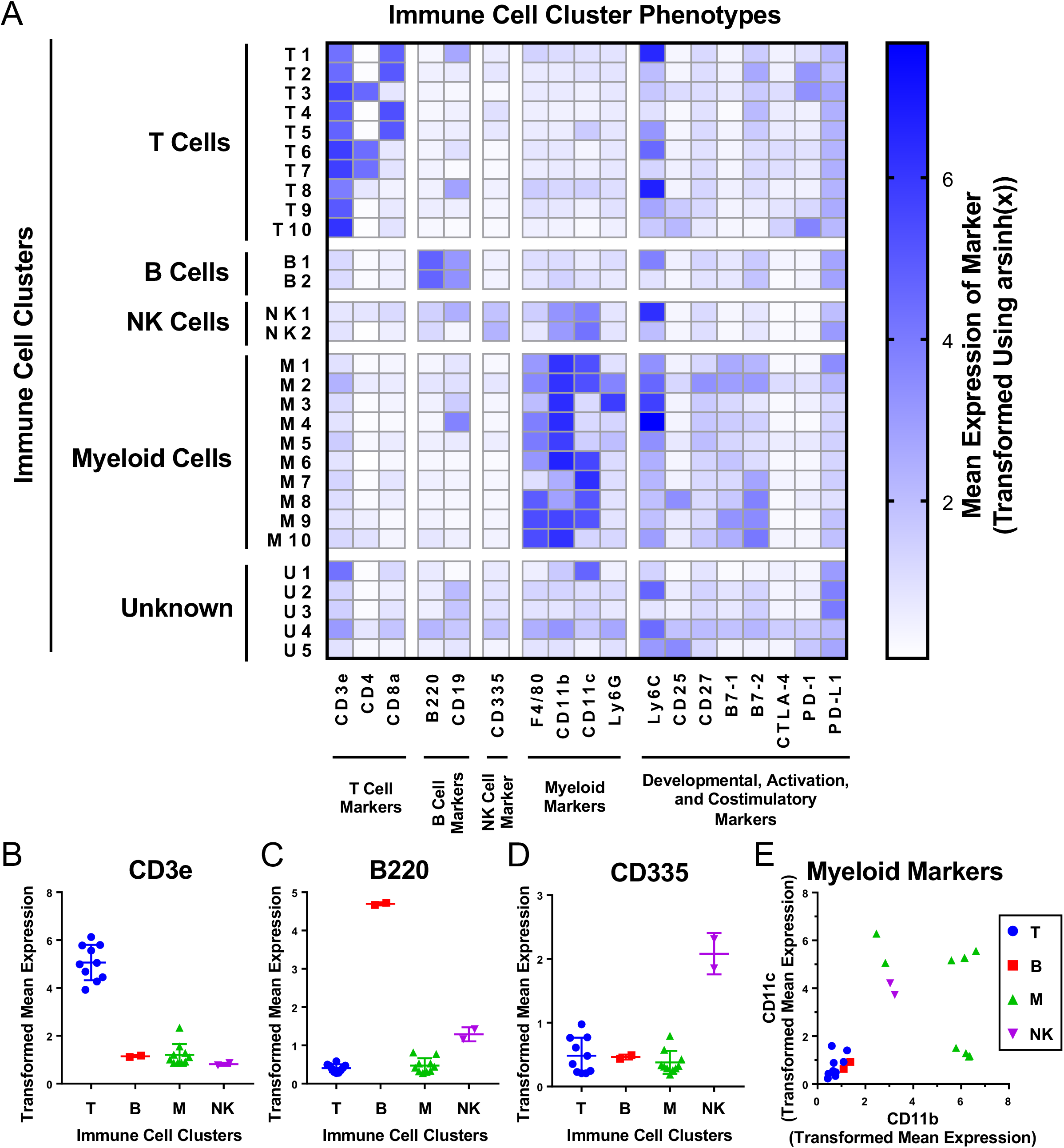
Unsupervised clustering of immune cells produces 29 phenotypically distinct clusters. (A) Heat map showing the phenotypes of the 29 immune cell clusters generated using X-shift clustering on immune cells detected from aging mouse prostates, bladders, and kidneys using CyTOF. Clusters were separated into T cells (CD3, CD4, CD8), B cells (B220, CD19), NK cells (CD335), myeloid cells (F4/80, CD11b, CD11c, and Ly6G) and unknown by expression of immune lineage markers. Shading represents mean marker expression transformed by arsinh(x). (B–E) Immune cell cluster expression of lineage markers for T cells (CD3) (B), B cells (B220) (C), NK cells (CD335) (D), and myeloid cells (CD11b and CD11c) (E). Data represents mean marker expression in each cluster transformed by arsinh(x) ± SD between clusters of the same broad immune cell type. Kruskal-Wallis, p < 0.001 (CD3), p < 0.05 (B220), p = 0.106 (CD335). Two-way ANOVA, p < 0.0001 (myeloid markers). See also Figures S2–S4.

We noticed 6 immune cell clusters with positive staining for both Ly6C and CD19 (T1, T8, B1, NK1, M4, U2) (Figure 2A). Since CD19 is only expected on B cells, expression on other cell types was thought to be unlikely (Engel et al., 1995). We plotted Ly6C against CD19 and found interference between the channels. At high levels of Ly6C, there was overflow of signal into the CD19 channel, resulting in a linearly proportional increase in signal between the two channels (Figure S3A). Ly6C was conjugated to neodymium-150, and CD19 was conjugated to erbium-166. These two lanthanide isotopes differ by 16 Da, consistent with known effects of oxidation of neodymium-150 by oxygen-16 as the sample is ionized to plasma in the mass cytometer (Figure S3B) (Leipold et al., 2015). We checked every other combination of the 19 markers used and found no other instances of signal spillover (Figure S3C). The effects of signal spillover from Ly6C into CD19 taken into account, high Ly6C expression may be causing T1, T8, NK1, M4, and U2 to appear as CD19^+^. However, expression of the B cell marker B220 by B1 indicates that its CD19 signal is likely true (Coffman and Weissman, 1981).

### Age-related changes in the mouse prostate immune compartment arise early in adulthood

Having used unsupervised clustering to subdivide immune cells in the mouse prostate into distinct subsets, we next looked at how these populations change throughout adulthood. Spearman correlation analysis was performed to determine whether each immune cell cluster was enriched (ρ > 0) or depleted (ρ < 0) with age. We found that 19/29 immune cell clusters were significantly correlated with age (ρ ≠ 0, P_ρ_ < 0.05). Consistent with the overall increase in T and B lymphocytes in the mouse prostate with age, 8/10 T cell clusters and 2/2 B cell clusters were significantly enriched with age. Similarly, we found that 1/2 NK cell clusters and 5/10 myeloid cell clusters had a significant negative correlation with age (Figure 3A and Table S1). There was an extreme outlier in the abundance of cluster M2 by one of the 9-month-old mice. When the outlier was excluded, we found a significant negative correlation (P_ρ_ < 0.05) between M2 abundance in the prostate and age (Figure S4).

**Figure 3.**
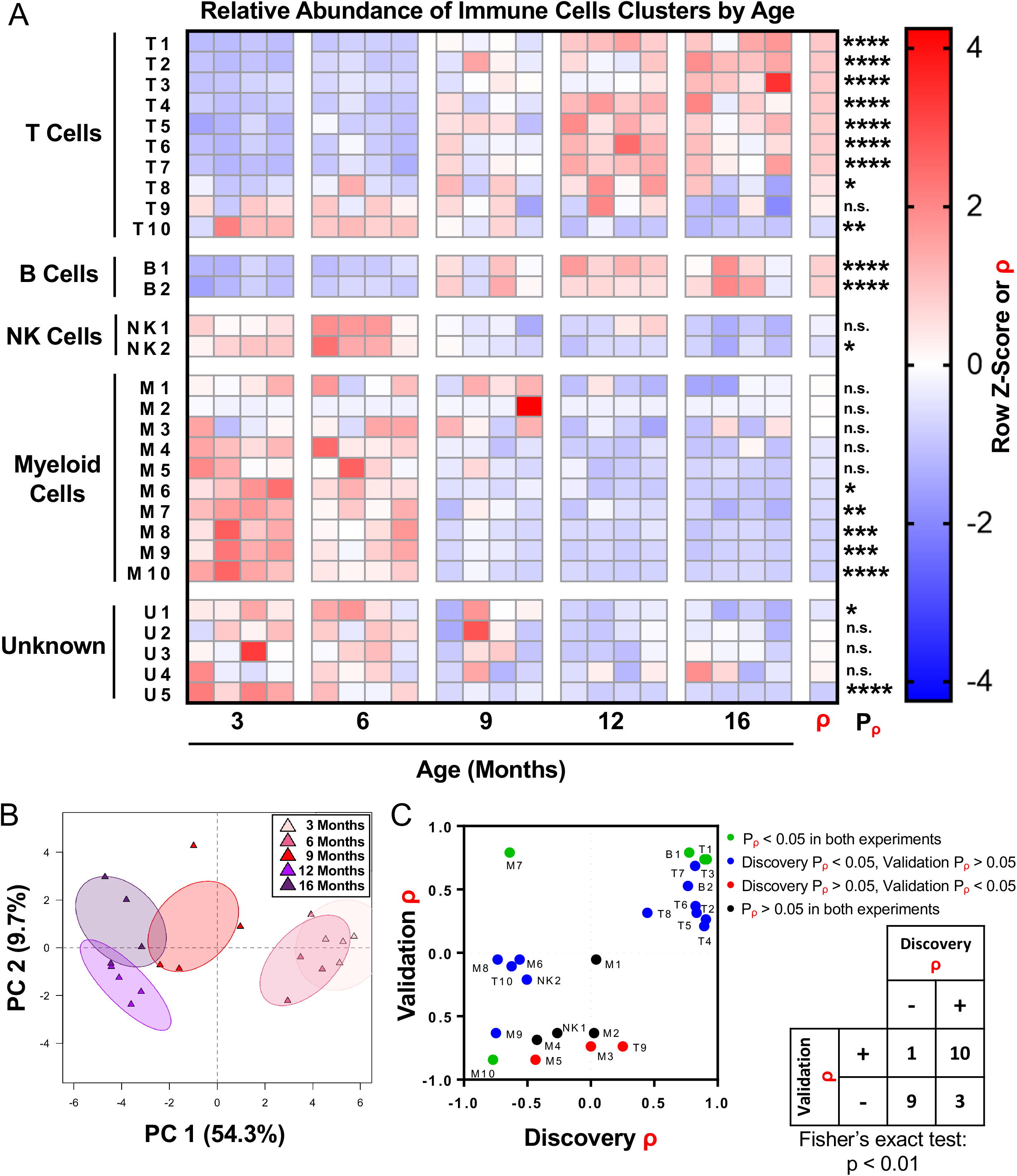
The adult mouse prostate immune microenvironment changes progressively with age. (A) Heat map showing changes to immune cell cluster abundance in the aging adult mouse prostate, correlation with age (ρ), and associated p-value (P_<_). Shading indicates abundance represented as a row z-score except where Spearman correlation (ρ) is indicated. Data represents 4 biological replicates at each age. *p < 0.05, **p < 0.01., ***p < 0.001, ****p < 0.0001. n.s., not significant, p ≥ 0.05. (B) Principal component analysis (PCA) was performed on immune cell cluster frequencies for mouse prostates at different ages. Ellipses represent 95% confidence intervals for 4 biological replicates at each age. PC, principal component. (C) Twenty-four clusters of known cell type were identified in a separate validation experiment with prostates from mice 4-, 9-, and 15-months-old (each n = 3), and correlation with age (ρ) and associated p-value (P_ρ_) were calculated. Left: Dot plot compares correlation coefficients for each immune cell cluster in the initial (discovery) and validation experiments. Coloring of points represents significance (P_ρ_). Right: Contingency table comparing the direction of correlation with age (ρ > 0 or ρ < 0) for identified immune cell clusters between the discovery and validation experiments. M3 was not counted because ρ = 0 in the discovery experiment. Fisher’s exact test was performed to evaluate the distribution of clusters in the four quadrants of the graph. See also Figure S5 and Tables S1–S3.

Principal component analysis was performed to assess the progression of changes to the mouse prostate immune microenvironment. Prostates from 3- and 6-month-old mice clustered together as did those from 12- and 16-month-old mice, demonstrating the similarity in immune microenvironments of prostates in these age groups (Figure 3B). Prostates from 9-month-old mice clustered between the other ages, demonstrating an intermediate immune phenotype at this age. These results show that there is a shift in the inflammatory microenvironment of the mouse prostate occurring between 6 and 12 months of age.

To validate these changes to the immune microenvironment of the mouse prostate, we performed a second CyTOF immunophenotyping experiment with a panel of the same markers and mice 4-, 9-, and 15-months-old (each n = 3) (Table S2). The 24 known immune cell clusters from the discovery experiment were matched to immune cells from the validation experiment, and Spearman correlation analysis was performed (Figure S5 and Table S3). Four of the 24 immune cell clusters (T1, T3, B1, and M10) were validated (same sign for ρ and P_ρ_ < 0.05 in both discovery and validation experiments) (Figure 3C). Fifteen of the 24 immune cell clusters had the same direction of correlation with age in both the discovery and validation experiments but failed to meet the barrier of statistical significance in both experiments. The majority of these were significant in the discovery but not the validation experiments, likely due to the low statistical power of n = 3 in the validation. Overall, there was a statistically significant correlation between the sign of ρ for immune cell clusters in each experiment (p < 0.01, Fisher’s exact test), supporting the reproducibility of our CyTOF immunophenotyping (Figure 3C).

Because CyTOF is not a widely available technique, we sought to provide a more accessible way to study these age-related immune microenvironment changes in the mouse prostate. Guided by the phenotypes of the validated immune cell clusters, a simpler gating scheme was constructed which can identify the 4 validated clusters (T1, T3, B1, and M10) and 5 additional clusters that were significant in the discovery experiment (T2, T7, T10, B2, and M9) using a reduced number of markers (Table 2 and Figure 4A). The reduced panel has fewer markers to be used with more widely accessible 12-color flow cytometry (De Rosa et al., 2003). Importantly, manual gating to reidentify these immune cell clusters in the discovery experiment produced populations with significant correlations with age (P_ρ_ < 0.0001) that are consistent with correlations found using X-Shift clustering (Figures 4B–4D).

**Figure 4.**
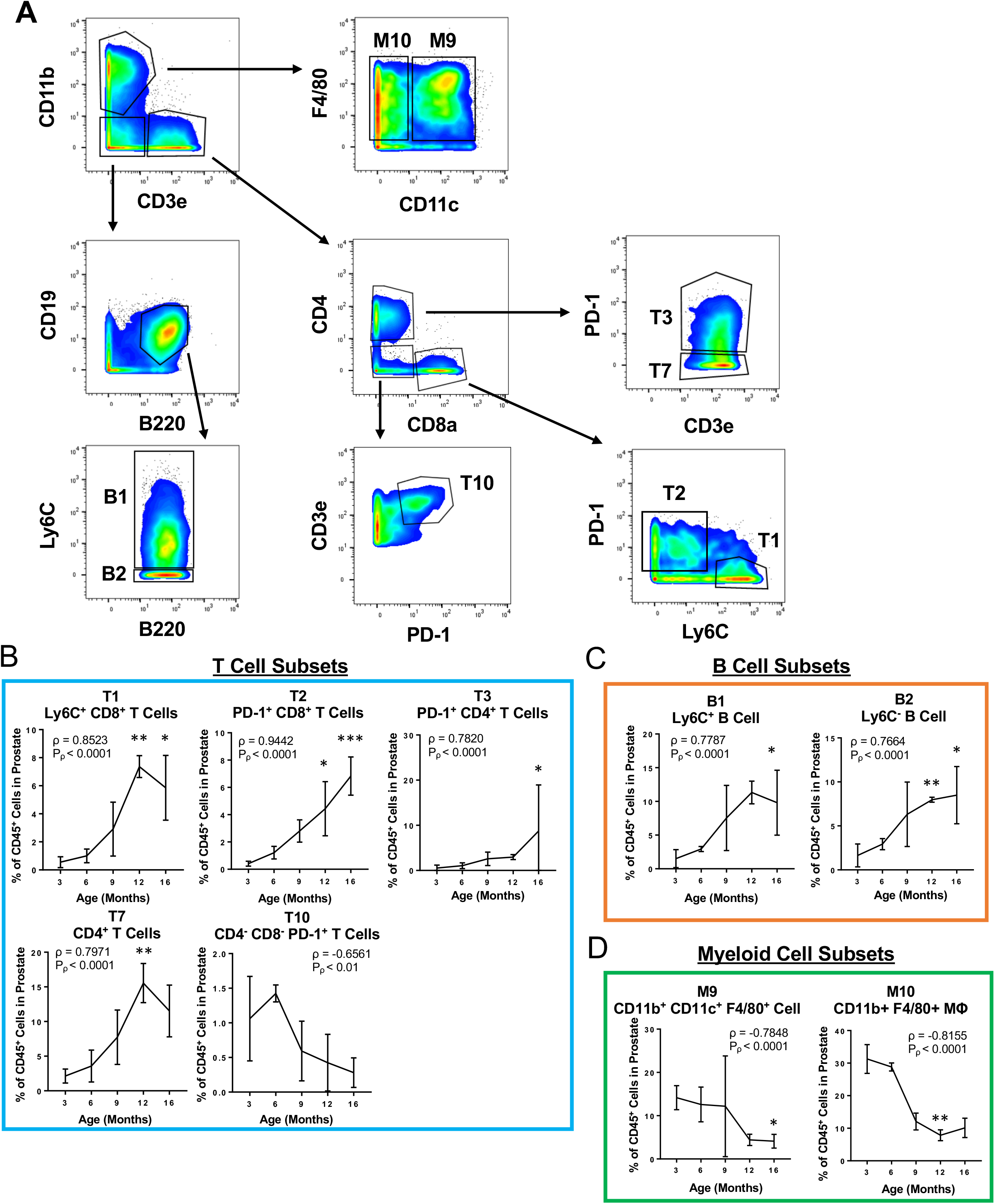
Simplified gating scheme to identify immune cell clusters that change in abundance with age in the mouse prostate. (A) Bivariate plots showing a simplified 11 marker panel (CD45, CD3e, CD4, CD8a, CD19, B220, F4/80, CD11b, CD11c, Ly6C and PD-1) used to identify 4 validated immune cell clusters (T1, T3, B1, and M10) and 5 others (T2, T7, T10, B2, and M9). (B–D) Quantification of immune cell subset abundance by proportion of total CD45^+^ cells in the mouse prostate identified using the simplified 11 marker panel separated into T cells (B), B cells (C), and myeloid cells (D). Data represents mean ± SD of 4 biological replicates at each age. MΦ, macrophage. Spearman correlation coefficient (ρ) and associated p-value (P_–_) represent the correlation with age. Kruskal-Wallis, p < 0.05 (T3, T10, B1, B2, M9), p < 0.01 (T1, T2, T7, M10). Dunn’s multiple comparisons test against 3-months-old, *p < 0.05, **p < 0.01, ***p < 0.001.

**Table 2.** Simplified panel for identifying immune cells in the mouse prostate that change with age.

| Marker |
| --- |
| CD45 |
| CD3e |
| CD4 |
| CD8a |
| F4/80 |
| CD11b |
| CD11c |
| B220 |
| CD19 |
| Ly6C |
| PD-1 |

### Mouse prostate and bladder share features of inflammaging

We performed CyTOF on bladders and kidneys from 3- and 16-month-old mice (each n = 4) to compare aging of the prostate to other urogenital tissues with age-related cancer risk (Shariat et al., 2010; Vogelzang and Stadler, 1998). Immune cells from these tissues were included for X-Shift clustering so that we could examine age-related changes to the same 29 immune cell clusters as the prostate (Table S1). In the mouse bladder, we identified significant correlations between age and 17/29 immune cell clusters (ρ ≠ 0, P_ρ_ < 0.05) (Figure S6A). The immune profiles of the 3- and 16-month-old mouse kidneys were similar, with only 5/29 immune cell clusters significantly correlated with age (Figure S6B). Principal component analysis was performed on the 3- and 16month old samples from the prostate, bladder, and kidney to assess the similarities in how different urogenital tissues age (Figure 5A). Three-month-old bladder and prostate clustered together as did 16-month-old bladder and prostate, suggesting these tissues are similar in how their immune microenvironments change with age. Both ages of kidney clustered together near the 16-month-old bladder, suggesting that the kidney immune microenvironment did not change with age and was most similar to the 16-month-old bladder.

**Figure 5.**
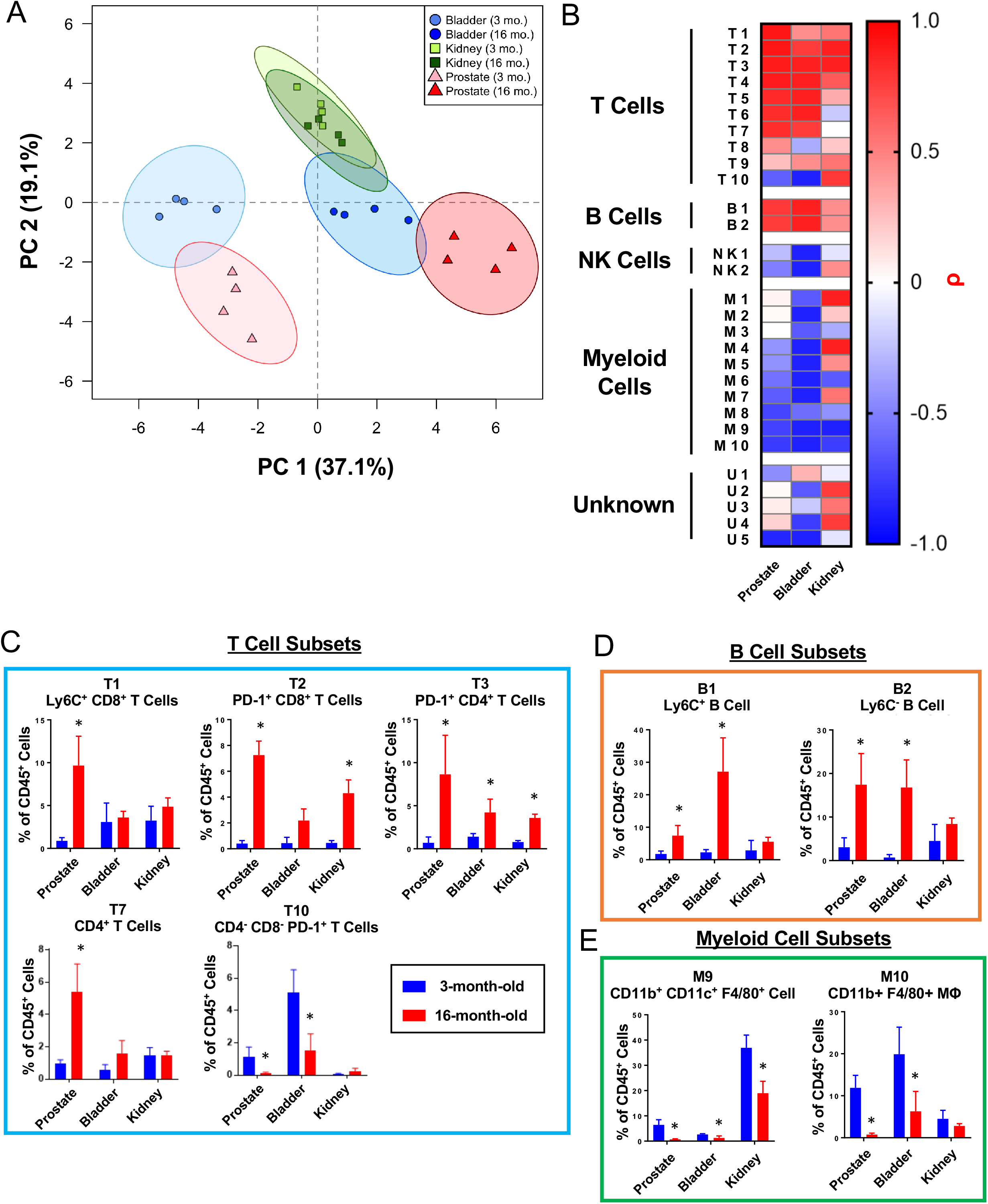
Comparison of age-related changes to the immune microenvironment of the mouse bladder, kidneys, and prostate. (A) Principal component analysis (PCA) was performed on immune cell cluster frequencies for mouse prostates, bladders, and kidneys at 3- and 16-months-old. Ellipses represent 95% confidence intervals for 4 biological replicates at each age and tissue. PC, principal component. (B) Heat map showing correlations with age (ρ) for immune cells clusters from mouse prostate, bladder, and kidneys irrespective of p-value. Shading represents Spearman correlation (ρ). (C–E) Quantification of immune cell cluster abundance at 3 and 16 months of age in mouse prostate, bladder, and kidney separated into T cells (C), B cells (D), and myeloid cells (E). Mann Whitney U test between 3- and 16-months-old, *p < 0.05. Data represents mean ± SD of 4 biological replicates at each age and tissue. See also Figure S6 and Table S1.

To better understand these relationships, we compared the correlation coefficients (ρ) for each immune cell cluster, irrespective of p-value, in the prostate, bladder, and kidney (Figure 5B). Thirteen of 29 clusters were correlated with age in the same direction in all three tissues (all ρ > 0 or ρ > 0) (T1-T4, T9, B1-B2, NK1, M6, M8-M10, and U5). Among the other immune cell clusters, prostate and bladder shared more similarities to each other than to kidney (Figure 5B–5E and Figure S6C). These results demonstrate that the immune compartments of mouse urogenital tissues age differently and that the aging prostate more closely resembles the aging bladder than the kidney.

## DISCUSSION

Inflammation in the prostate and aging are both associated with an increased risk for developing prostate cancer (Gurel et al., 2014; Sfanos and De Marzo, 2012). Our group and others have previously reported that the mouse prostate experiences an increase in immune cells as it ages, however the dynamics of how these changes occur were unclear (Bianchi-Frias et al., 2010; Crowell et al., 2019). In this study, we characterized how the inflammatory microenvironment of the adult mouse prostate changes during aging using highly-multiplexed single-cell mass cytometry. This dataset presents the most comprehensive profiling of the aging adult mouse prostate immune profile to date.

By 3 months of age, male mice are sexually mature and the prostate has completed development (Marker et al., 2003). At this age, myeloid cells make up the majority of immune cells in the mouse prostate (Figure 1E). As mice age, the immune compartment of the mouse prostate undergoes an expansion of T and B lymphocytes resulting in a profound shift in the tissue’s immunological milieu (Figures 1C and 1D). Importantly, this age-related expansion of lymphocytes in the prostate happens in contrast to the age-related myeloid bias of hematopoiesis (Mann et al., 2018). As mice age, different processes including bone marrow stem cell niche remodeling, inflammatory cytokines, and telomere dysfunction skew hematopoiesis towards myeloid lineages (Ergen et al., 2012; Ho et al., 2019; Ju et al., 2007). Aging is also associated with involution of the thymus, resulting in reduced production of naïve T cells (Palmer, 2013). Additionally, B cell development is impaired in aged mouse bone marrow (Zharhary, 1988). These many processes converge to reduce lymphocyte output and increase myeloid differentiation. The contrasting lymphocyte expansion in the mouse prostate suggests that the prostate contains a somewhat unique aging immune microenvironment.

We used 19 surface markers to identify 29 phenotypically-distinct clusters of immune cells in the mouse prostate (Figure 2). We then tracked how the abundance of each immune cell cluster changes during adult mouse aging by comparing tissue aged 3-, 6-, 9-, 12-, and 16-months. We found that there is a progressive change in the immune microenvironment of the mouse prostate taking place between 6 and 12 months of age (Figures 3A and 3B). A separate validation experiment demonstrated that the majority of age-related changes to specific immune cell clusters were reproducible (Figure 3C). Future studies will be necessary to determine the mechanisms promoting an altered prostate immune compartment between 6 and 12 months of age.

Our findings reveal that aging of the mouse prostate immune compartment is complex. In the aging mouse prostate, the proportion of total T and B cells increases as the proportion of total NK and myeloid cells decreases (Figures 1C–1E). Analyzing these changes at the more granular level of immune cell cluster shows greater complexity in these changes to the immune microenvironment. While all B cell clusters were enriched with age, we identified T cell clusters which did not change or were reduced in abundance with age. Similarly, while the majority of NK and myeloid cell clusters decreased with age, we found NK and myeloid cell clusters which did not significantly change in proportion with age (Figure 3A). These findings clarify that a subset of immune cells clusters accounts for the overall increase in T and B lymphocytes and decrease in NK and myeloid cells with age.

Though we used highly-multiplexed CyTOF to study changes to the prostate immune microenvironment, we sought to devise a simplified panel that could be accessible with more widely available technologies. We took the most reproducible immune cell cluster changes and designed a panel of 11 markers which can reidentify the same immune cell clusters that are significantly correlated with age (Table 2 and Figure 4). This reduced panel could be used to study changes to the mouse prostate immune microenvironment using more accessible 12-color flow cytometry (De Rosa et al., 2003).

Finally, we used CyTOF to survey the immune microenvironment of the aging mouse bladder and kidney to determine how the aging phenotype of the prostate compares to other urogenital tissues. Between the bladder and kidney, we found that the aging prostate’s immune compartment is more similar to the bladder (Figure 5 and Figure S6). The proximity of the prostate and bladder may explain their similar aging immune profiles. Both tissues are derived from the urogenital sinus, whereas the kidney develops from the ureteric bud and metanephric mesenchyme (McMahon, 2016; Staack et al., 2003). The similarity in immune compartment aging between mouse prostate and bladder may also be reflective of their shared developmental origin. Understanding what drives similar age-related changes to the immune compartment in prostate and bladder nay reveal common approaches to prevent inflammaging and reduce age-related disease risk.

## LIMITATIONS OF THE STUDY

In this study, we evaluated immune cells in the aging mouse prostate using mass cytometry to take advantage of its high-throughput single-cell data. This approach requires dissociation of the tissue to a single-cell suspension using mechanical and enzymatic means. As a result, we were only able to profile immune cells that could survive the dissociation protocol. Between different organs and ages, we assume that specific immune cell types are equally likely to survive the dissociation protocol, allowing for relative comparisons to still be made. Other approaches that preserve tissue architecture, like immunohistochemistry (IHC), immunofluorescence (IF), and imaging mass cytometry (IMC), may be necessary to detect cell types that may not survive long enough for CyTOF detection.

For both weight and % CD45^+^ immune cells, there were time points whose mean values were greater than those at 3-months-old yet were not statistically significant (Figures S1A and S1B). The relatively low statistical power of Dunn’s test may be responsible for some intermediate ages not reaching the threshold of significance (Kromrey and La Rocca, 1995). Additional replicates would be useful in increasing the statistical power and determining whether these changes happen even earlier in adulthood than 12 months of age.

## RESOURCE AVAILABILITY

**Lead Contact**

Further information and requests for resources and reagents should be directed to and will be fulfilled by the Lead Contact, Andrew S. Goldstein.

## Materials Availability

This study did not generate new unique reagents.

## Data and Code Availability

The mass cytometry data supporting the current study has not been deposited in a public repository but is available from the corresponding author upon request.

## ACKNOWLEDGMENTS

Special thanks to Miriam Guemes for CyTOF sample acquisition and Johnny Diaz for sample preparation. Mass cytometry was performed in the UCLA Jonsson Comprehensive Cancer Center (JCCC) and Center for AIDS Research Flow Cytometry Core Facility which is supported by NIH awards P30 CA016042 and 5P30 AI028697. The purchase of the Helios mass cytometer that was used in this work was, in part, supported by funds provided by the James B. Pendleton Charitable Trust. J.J.F. was supported by scholarships from the UCLA Minor in Biomedical Research and the Silva Endowment as part of the Undergraduate Research Scholars Program at UCLA. H.I.N. was supported by the Eugene V. Cota-Robles Fellowship. A.S.G. is supported by the Spitzer Family Foundation Fund and the Gill Endowment. This work was supported by the American Cancer Society (RSG-17-068-01-TBG), U.S. Department of Defense (W81XWH-13-1-0470), STOP CANCER, NIH/NCI (R01CA237191 and P50CA092131/UCLA SPORE in Prostate Cancer), UCLA Eli and Edythe Broad Center of Regenerative Medicine and Stem Cell Research Rose Hills Foundation Innovator Grant, and support from UCLA’s Jonsson Comprehensive Cancer Center, Broad Stem Cell Research Center, Clinical and Translational Science Institute, and the Institute of Urologic Oncology. We also thank UCLA’s Institute for Quantitative and Computational Biology.

## AUTHOR CONTRIBUTIONS

J.J.F., T.H., H.I.N., and A.S.G. conducted the experiments. J.J.F., H.I.N., and A.S.G. designed the experiments. J.J.F. analyzed flow cytometry and mass cytometry data. J.J.F. and A.S.G. wrote and edited the manuscript. A.J.G. assisted with CyTOF samples and wrote part of the manuscript. B.L.S. performed principal component analysis and wrote part of the manuscript. A.S.G. procured funding and supervised the experiments.

## EXCEL TABLE TITLES

**Table S1. Abundance of X-Shift-generated immune cell clusters in the aging mouse prostate, bladder, and kidney from the discovery experiment. Related to Figures 3 and 5.**

**Table S3. Abundance of phenotypically matched immune cell clusters in the aging mouse prostate from the validation experiment. Related to Figure 3.**

