## Supplemental files for "Highly multiplexed immune profiling throughout adulthood reveals kinetics of lymphocyte infiltration in the aging mouse prostate"

### SUPPLEMENTAL INFORMATION

#### SUPPLEMENTAL FIGURES

### Figure S1.

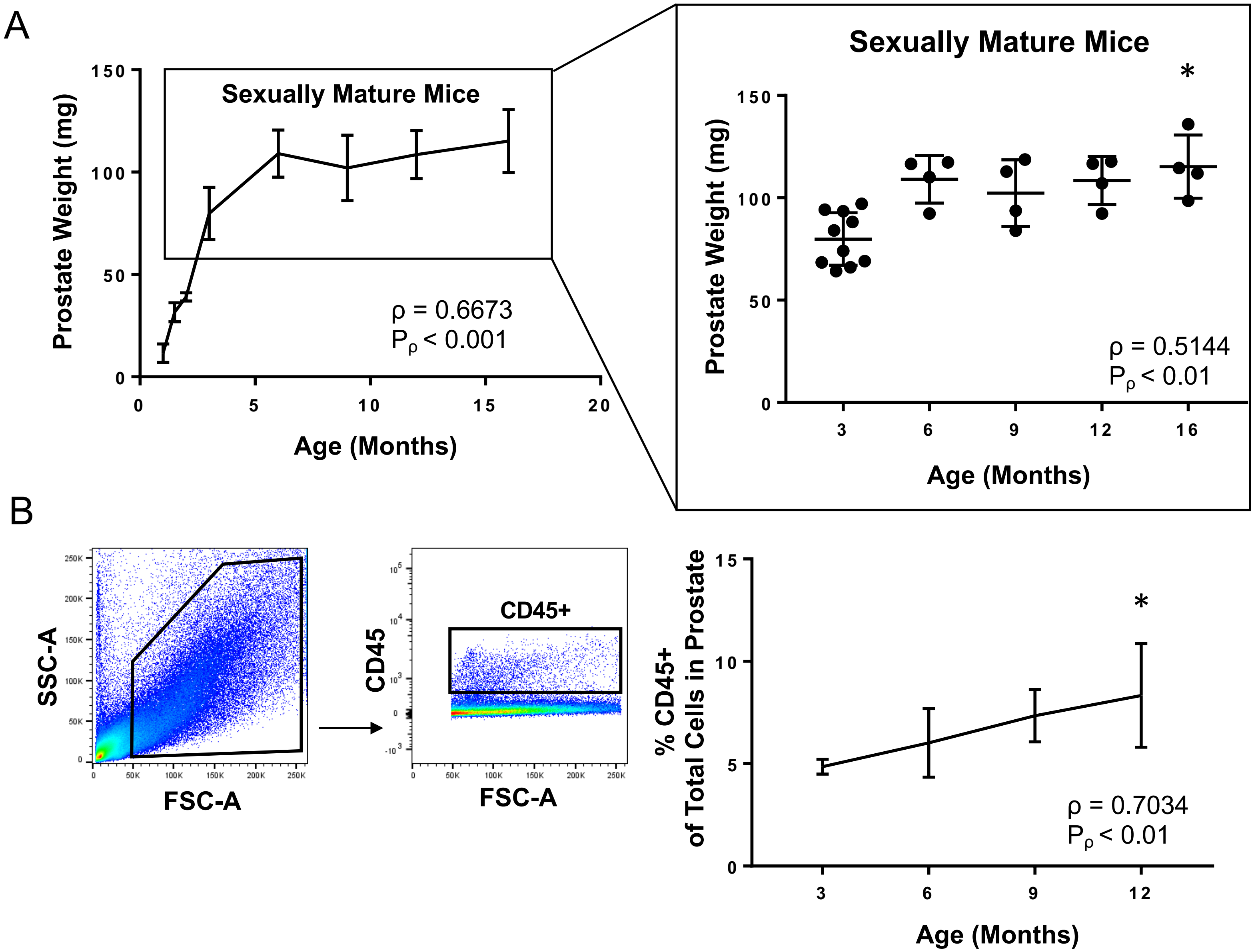
A
Sexually Mature Mice
Prostate Weight (mg)
Age (Months)
ρ = 0.6673
$P_\rho$ < 0.001
Sexually Mature Mice
*
Prostate Weight (mg)
Age (Months)
ρ = 0.5144
$P_\rho$ < 0.01
B
SSC-A
FSC-A
CD45+
CD45
FSC-A
% CD45+
of Total Cells in Prostate
Age (Months)
*
ρ = 0.7034
$P_\rho$ < 0.01

**Figure S1. Increased weight and inflammation in the aging adult mouse prostate. Related to Figure 1.**

(A) Wet weight of prostates from mice of different ages. Left: Prostate weights of mice aged 1–16 months. Right: Prostate weights of sexually mature mice aged 3–16 months. Data represents mean ± SD of 2–10 biological replicates at each age. Spearman correlation coefficient (ρ) and associated p-value ($P_\rho$) represent the correlation with age. Kruskal-Wallis, $p < 0.05$ (sexually mature mice). Dunn's multiple comparisons test against 3-month-old mice, *$p < 0.05$.

(B) Analysis of immune cells in the prostate by flow cytometry. Left: Gating scheme to identify immune cells ($CD45^+$). FSC-A, forward scatter area; SSC-A, side scatter area. Right: Quantification of $CD45^+$ immune cell frequency in the mouse prostate. Data represents mean ± SD of 4 biological replicates at each age. Spearman correlation coefficient (ρ) and associated p-value ($P_\rho$) represent the correlation with age. Kruskal-Wallis, $p < 0.05$. Dunn's multiple comparisons test against 3-month-old mice, *$p < 0.05$.

### Figure S2.

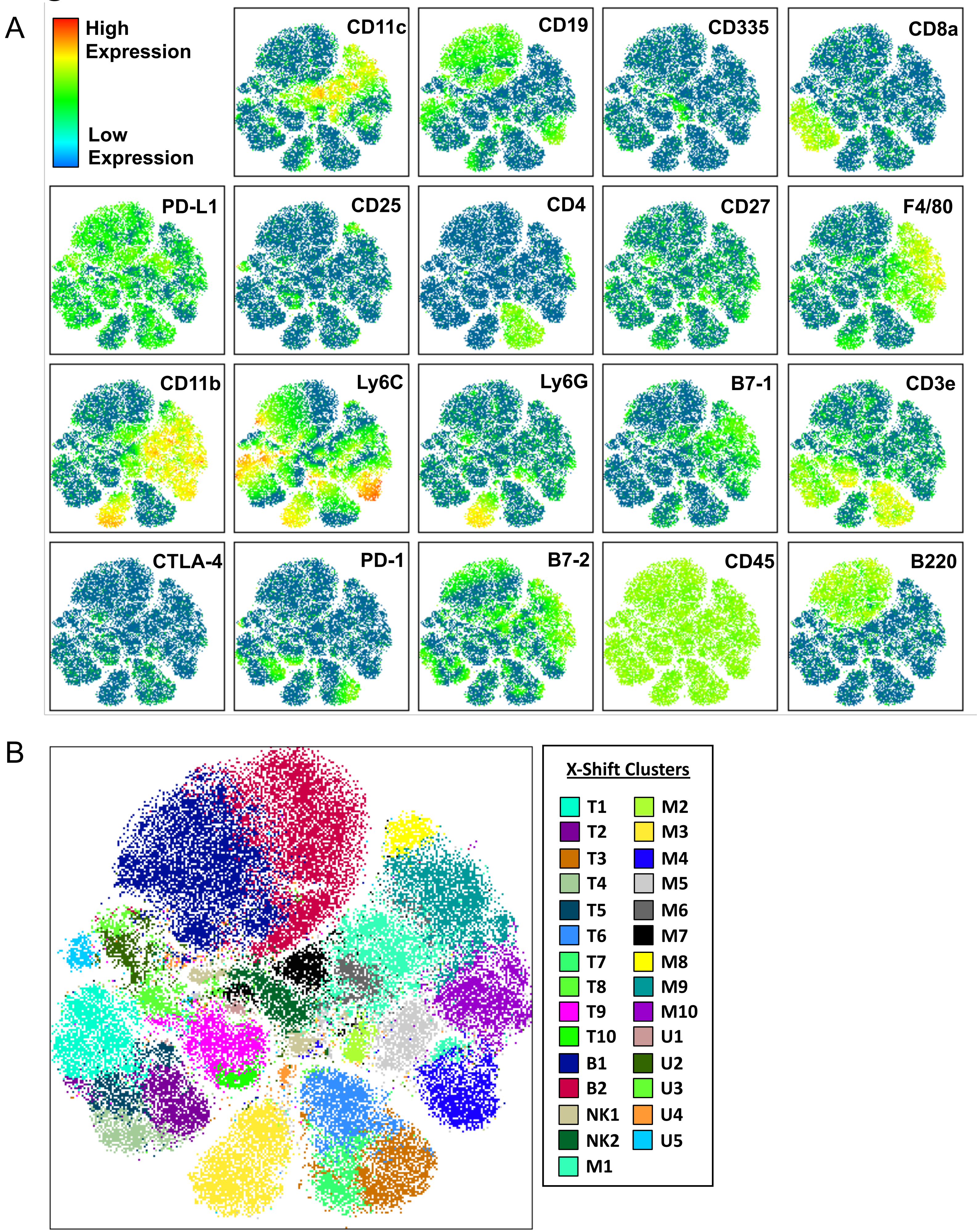
A
High
Expression
Low
Expression
CD11c
CD19
CD335
CD8a
PD-L1
CD25
CD4
CD27
F4/80
CD11b
Ly6C
Ly6G
B7-1
CD3e
CTLA-4
PD-1
B7-2
CD45
B220
B
X-Shift Clusters
T1
T2
T3
T4
T5
T6
T7
T8
T9
T10
B1
B2
NK1
NK2
M1
M2
M3
M4
M5
M6
M7
M8
M9
M10
U1
U2
U3
U4
U5

**Figure S2. Visualizing the heterogeneity of immune cells in mouse urogenital tissues with dimensional reduction and clustering algorithms. Related to Figures 1 and 2.**

(A) t-SNE plot generated from the immune cells from mouse prostate, bladder, and kidney. Color mapping shows expression of each of the 19 markers in the CyTOF panel.

(B) t-SNE plot showing 29 X-Shift generated immune cell clusters.

### Figure S3.

A

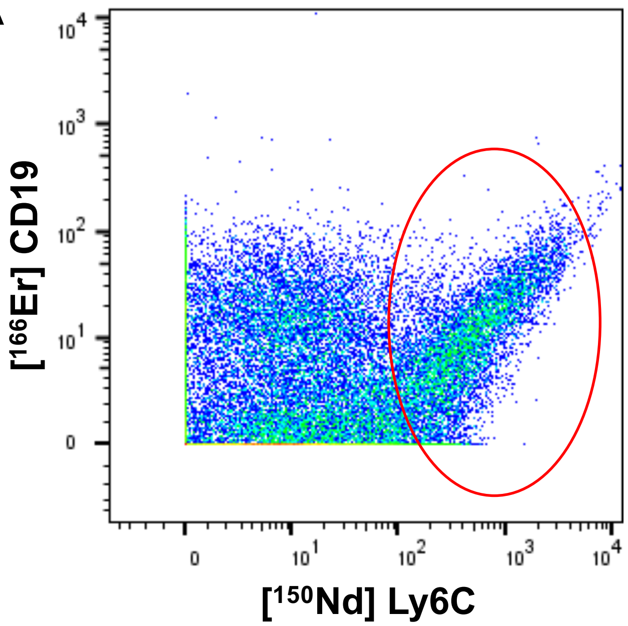


B

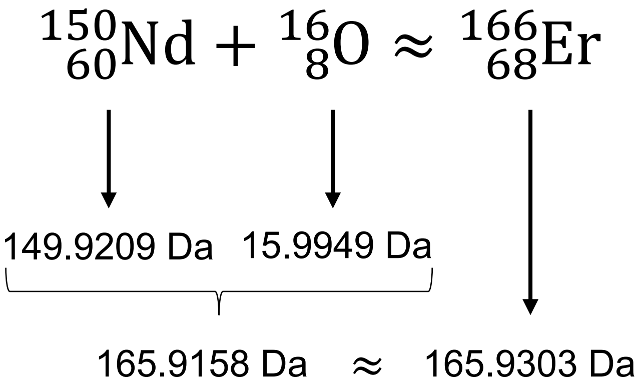


C

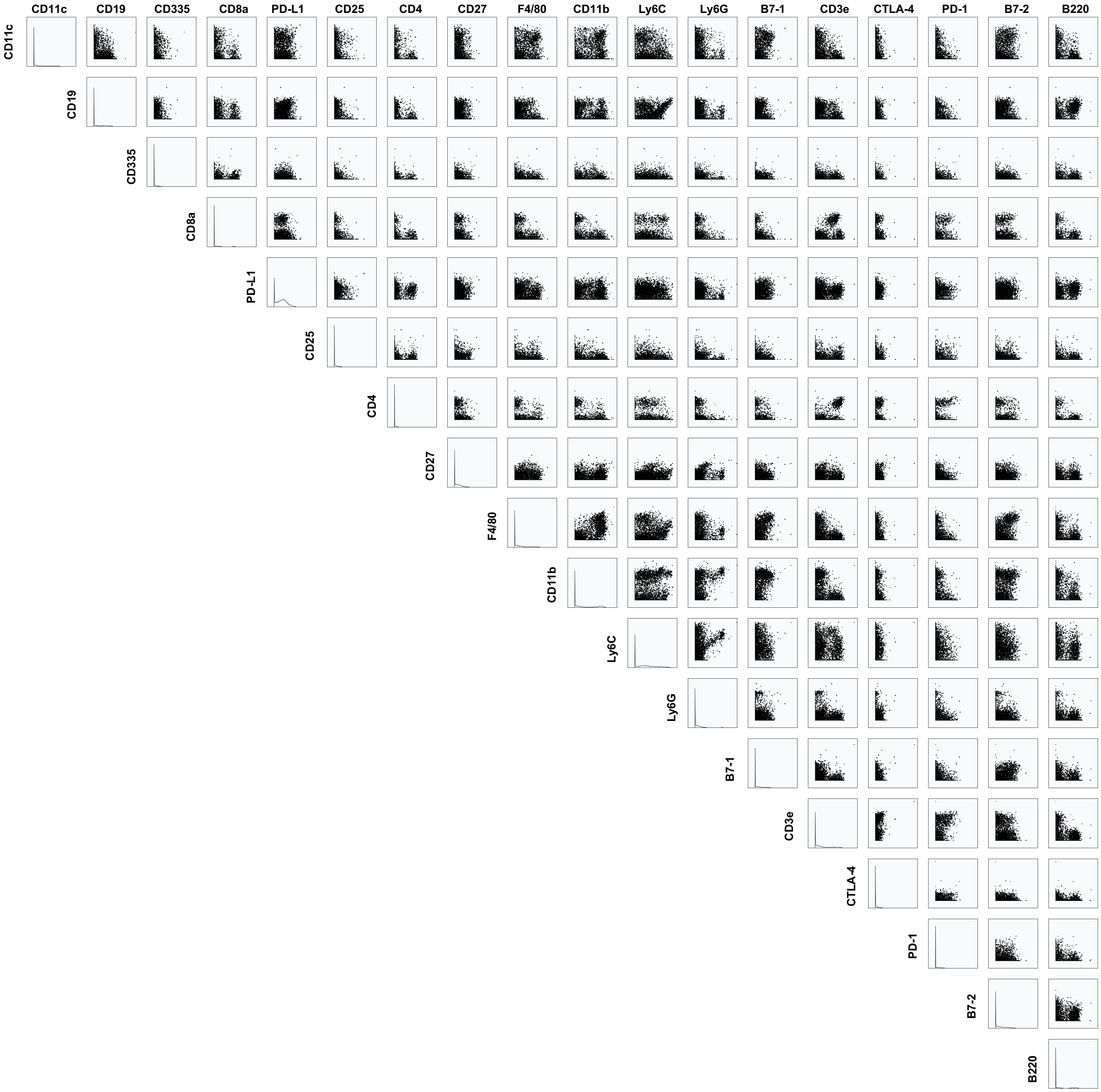

**Figure S3. Oxidation of the neodymium-150 ion leads to spillover into the CD19 channel. Related to Figure 2.**

(A) Bivariate plot between [$^{150}$Nd] Ly6C and [$^{166}$Er] CD19 CyTOF channels showing all CD45$^{+}$ cells from the discovery experiment. The red ellipse identifies the area on the plot in which there is a proportional increase in both channels starting only at high levels of [$^{150}$Nd] Ly6C signal.

(B) When neodymium-150 is oxidized by oxygen-16, the most abundant isotope of oxygen, the resulting molecule contains 68 protons and 98 neutrons for a total mass of 166 Da. When detected by the time-of-flight mass spectrometer, the oxidized neodymium-150 has approximately the same apparent mass as erbium-166, causing spillover of the [$^{150}$Nd] Ly6C signal into [$^{166}$Er] CD19 when Ly6C is very highly expressed. Masses of isotopes were retrieved from PubChem (Kim et al., 2019).

(C) N x N plot showing all possible bivariate comparisons between the 19 CyTOF markers used in the discovery experiment. Boxes comparing a marker to itself show histograms of that marker's expression by immune cells in the discovery experiment.

### Figure S4.

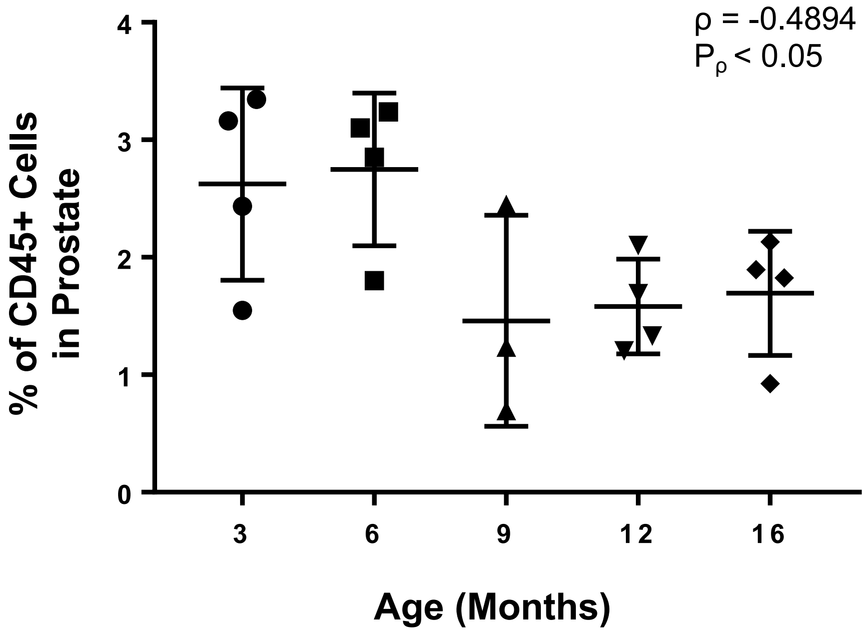
ρ = -0.4894
$P_\rho < 0.05$
% of CD45+ Cells in Prostate
4
3
2
1
0
3
6
9
12
16
Age (Months)

**Figure S4. Analysis of M2 frequency in mouse prostate without an outlier. Related to Figure 2.**

Quantification of % M2 of $CD45^+$ cells in the aging mouse prostate without an outlier at 9-months-old. Spearman correlation coefficient (ρ) and associated p-value ($P_\rho$) represent the correlation with age. Data represents mean ± SD of 3–4 biological replicates at each age.

### Figure S5.

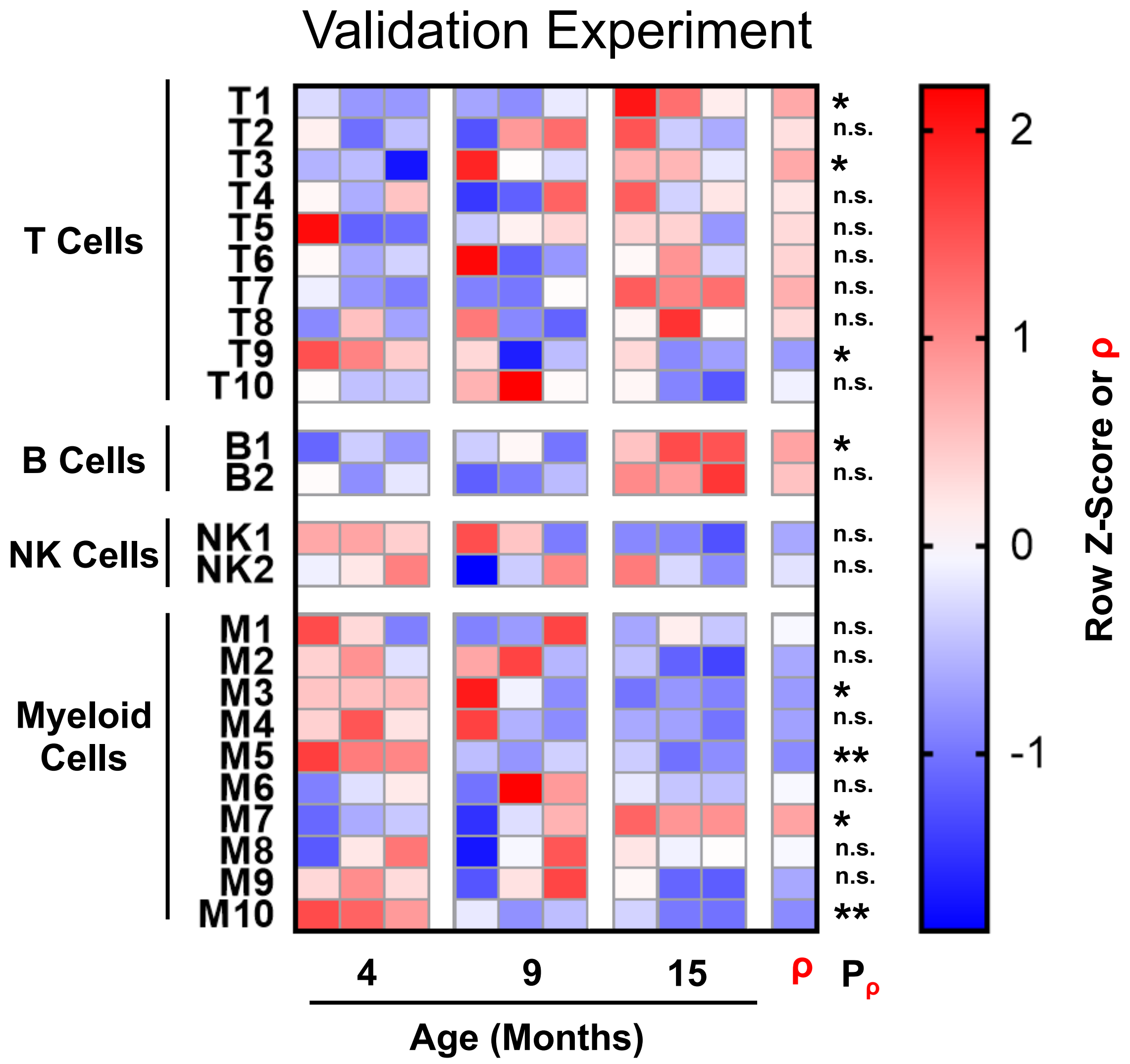

Validation Experiment
T Cells
T1
T2
T3
T4
T5
T6
T7
T8
T9
T10
B Cells
B1
B2
NK Cells
NK1
NK2
Myeloid Cells
M1
M2
M3
M4
M5
M6
M7
M8
M9
M10
*
n.s.
*
n.s.
n.s.
n.s.
n.s.
n.s.
*
n.s.
*
n.s.
n.s.
n.s.
n.s.
n.s.
*
n.s.
**
n.s.
*
n.s.
n.s.
**
4
9
15
ρ
Pρ
Age (Months)
2
1
0
-1
Row Z-Score or ρ

**Figure S5. Age-related immune cell cluster enrichment in the prostates of mice in the validation experiment. Related to Figure 3.**

A separate validation experiment was performed, using CyTOF to profile immune cells in the prostates of mice ages 4-, 9-, and 15-months-old with a panel containing the same markers as the discovery experiment. Heat map shows changes to matched immune cell cluster abundance in the aging adult mouse prostate, correlation with age ($\rho$), and associated p-value ($P_\rho$). Shading indicates abundance represented as a row z-score except where Spearman correlation ($\rho$) is indicated. Data represents 3 biological replicates at each age. *$p < 0.05$, **$p < 0.01$. n.s., not significant, $p \geq 0.05$.

### Figure S6.

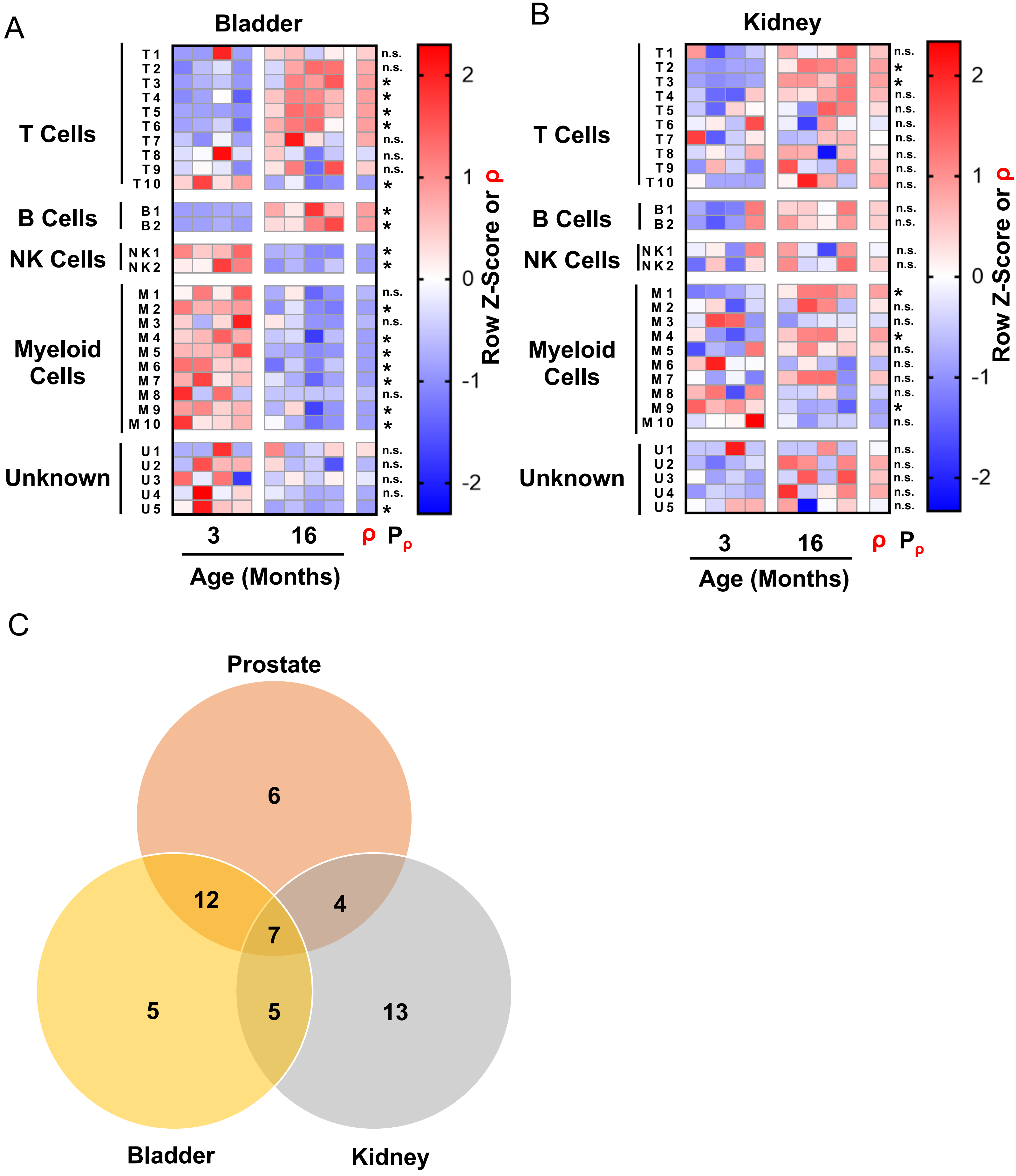
A
Bladder
T Cells
T 1
T 2
T 3
T 4
T 5
T 6
T 7
T 8
T 9
T 10
B Cells
B 1
B 2
NK Cells
N K 1
N K 2
Myeloid Cells
M 1
M 2
M 3
M 4
M 5
M 6
M 7
M 8
M 9
M 10
Unknown
U 1
U 2
U 3
U 4
U 5
n.s.
*
3
16
ρ
$P_\rho$
Age (Months)
Row Z-Score or ρ
2
1
0
-1
-2
B
Kidney
C
Prostate
Bladder
Kidney
6
12
4
7
5
5
13

**Figure S6. Mouse prostate and bladder share more age-related changes to their immune microenvironment with each other than to kidney. Related to Figure 5.**

(A–B) Heat maps showing changes to immune cell cluster abundance in the aging adult mouse bladder (A) and kidneys (B), correlation with age (ρ), and associated p-value ($P_\rho$). Shading indicates abundance represented as a row z-score except where Spearman correlation (ρ) is indicated. Data represents 4 biological replicates at each age. *$p < 0.05$. n.s., not significant, $p \geq 0.05$.

(C) Venn diagram showing the age-related immune cluster changes shared between the mouse prostate, bladder, and kidneys. An age-related immune cell cluster change is considered shared if each tissue has the same sign for the correlation with age (ρ both positive or negative) and if both are significant ($P_\rho < 0.05$). Clusters with no significant correlation with age in multiple tissues are also considered shared.

#### SUPPLEMENTAL TABLES

**Table S1. Abundance of X-Shift-generated immune cell clusters in the aging mouse prostate, bladder, and kidney from the discovery experiment. Related to Figures 3 and 5.**

See Excel Table S1.

**Table S2. CyTOF immunophenotyping antibody panel for validation experiment. Related to Figure 3.**

| Label | Target | Clone | Conjugation | Source |
|---|---|---|---|---|
| $^{89}Y$ | CD45 | 30-F11 | Pre-conjugated | Fluidigm |
| $^{139}La$ | CD27 | LG.3A10 | Maxpar kit | BioLegend |
| $^{143}Nd$ | CD86 (B7-2) | GL1 | Maxpar kit | BioLegend |
| $^{146}Nd$ | F4/80 | BM8 | Pre-conjugated | Fluidigm |
| $^{147}Sm$ | CD80 (B7-1) | 16-10A1 | Maxpar kit | BioLegend |
| $^{148}Nd$ | CD11b | M1/70 | Pre-conjugated | Fluidigm |
| $^{150}Nd$ | Ly6C | HK1.4 | Pre-conjugated | Fluidigm |
| $^{151}Eu$ | Ly6G | 1A8 | Pre-conjugated | Fluidigm |
| $^{152}Sm$ | CD3e | 145-2C11 | Pre-conjugated | Fluidigm |
| $^{153}Eu$ | CD335 (NKp46) | 29A1.4 | Pre-conjugated | Fluidigm |
| $^{154}Sm$ | CD152 (CTLA-4) | UC10-4B9 | Pre-conjugated | Fluidigm |
| $^{155}Gd$ | CD25 | 3C7 | Maxpar kit | BioLegend |
| $^{156}Gd$ | CD4 | RM4-5 | Maxpar kit | BioLegend |
| $^{159}Tb$ | CD279 (PD-1) | 29F.1A12 | Maxpar kit | BioLegend |
| $^{166}Er$ | CD19 | 6D5 | Pre-conjugated | Fluidigm |
| $^{168}Er$ | CD8a | 53-6.7 | Pre-conjugated | Fluidigm |
| $^{169}Tm$ | CD274 (PD-L1) | 10F.9G2 | Maxpar kit | BioLegend |
| $^{176}Yb$ | CD45R (B220) | RA3-6B2 | Pre-conjugated | Fluidigm |
| $^{209}Bi$ | CD11c | N418 | Pre-conjugated | Fluidigm |

**Table S3. Abundance of phenotypically matched immune cell clusters in the aging mouse prostate from the validation experiment. Related to Figure 3.**

See Excel Table S3.

#### TRANSPARENT METHODS

##### Animal Work

Male C57BL/6J and C57BL/6N mice from Jackson Laboratories and the UCLA Department of Radiation Oncology's animal core facility were used. The UCLA Division of Laboratory Animal Medicine (DLAM) bred and maintained the mice. The UCLA Animal Research Committee (ARC) and DLAM approved all animal protocols.

##### Tissue Isolation and Dissociation to Single Cells

Male mice of different ages were euthanized using carbon dioxide asphyxiation, then their urogenital tract and right kidneys were removed. Under a microscope, microdissection was performed to isolate the prostate and bladder from the urogenital tract. Tissues were mechanically dissociated with a razor blade then enzymatically dissociated in RPMI 1640 (Gibco) containing 10% fetal bovine serum (FBS), 1mg/mL collagenase type I (Gibco), and 0.1 mg/mL DNase (Sigma) at 37°C on a nutating platform for 60–90 minutes. After washing cell pellets with 1X DPBS (Gibco), cells were resuspended in 37°C TrypLE Express Enzyme, no phenol red (Thermo Fisher Scientific) for 5 minutes before being quenched by adding RPMI 1640. Cells were further disrupted using the shear stress generated by drawing through an 18G syringe. Finally, cells were sequentially passed through 100 μm and 70 μm cell strainers (Corning) to produce a single-cell suspension.

##### Flow Cytometry

Single-cell suspensions of $5 \times 10^4$ – $1 \times 10^6$ cells from mouse prostate tissue of different ages (each n = 4) were stained in cell staining buffer comprised of 1X DPBS (Gibco) with 5 g/L protease-free bovine serum albumin (Sigma-Aldrich) and 200 mg/L sodium azide (Sigma-Aldrich). Nonspecific antibody binding through Fc receptors was blocked using TruStain fcX (anti-mouse CD16/32) Antibody (BioLegend) according to the manufacturer's protocol. Cells were stained with rat anti-CD49f-PE (BioLegend), rat anti-CD326 (EpCAM)-APC/Cy7 (BioLegend), goat anti-Trop2-APC (R & D Systems), and rat anti-CD45-FITC (BioLegend) for 30 minutes at room temperature. Cells were washed with cell staining buffer then fixed in 1% paraformaldehyde (Electron Microscopy Sciences) for 10-15 minutes at 37°C. After fixation, cells were chilled on ice for 1 minute, washed with cell staining buffer, and stored at 4°C before analysis on a FACSCanto flow cytometer (BD Biosciences).

##### Antibodies for Mass Cytometry

Antibody panels were designed using the Maxpar Panel Designer software (Fluidigm) to check for interference between the channels. The discovery and validation experiments used the same list of markers but differed in the metal tags conjugated to the antibodies (Table 1 and Table S2). Antibodies were purchased pre-conjugated from Fluidigm or conjugated at the UCLA Jonsson Comprehensive Cancer Center (JCCC) and Center for

AIDS Research Flow Cytometry Core Facility using the Maxpar X8 Polymer Chemistry kit (Fluidigm) following the manufacturer's instructions. Antibody cocktails were prepared by adding 1.1 µL/sample of the conjugated antibody stock to a single tube then diluting to a volume of 50 µL/sample with cell staining buffer comprised of 1X DPBS (Gibco) with 5 g/L protease-free bovine serum albumin (Sigma-Aldrich) and 200 mg/L sodium azide (Sigma-Aldrich).

**Cell Surface Staining for Mass Cytometry**

Single-cell suspensions of $3 \times 10^5 – 1.8 \times 10^6$ cells from mouse prostate, bladder, and kidney tissue of different ages were stained in cell staining buffer comprised of 1X DPBS (Gibco) with 5 g/L protease-free bovine serum albumin (Sigma-Aldrich) and 200 mg/L sodium azide (Sigma-Aldrich). Staining was performed in a 96-well V-bottom plate. The discovery experiment was performed using prostates from 3-, 6-, 9-, 12-, and 16-months-old mice; bladders from 3- and 16-months-old mice; and kidneys from 3- and 16-months-old mice (each n = 4). The validation experiment was performed using prostates from mice aged 4-, 9-, and 15-months old (each n = 3). To distinguish between live and dead cells, cells were washed with cell staining buffer, resuspended in 200 µL cell staining buffer containing 1 µM Cell-ID Intercalator-$^{103}$Rh (Fluidigm), and incubated for 15 minutes at 37°C. Staining with rhodium-103 was quenched with 2 mL cell staining buffer. Next, cells were centrifuged at 400 x g for 5 minutes, resuspended in 50 µL cell staining buffer containing 1 µg/mL TruStain fcX (anti-mouse CD16/32) Antibody (BioLegend), and incubated at room temperature for 10 minutes. Next, 50 µL of the diluted antibody cocktail was added to each sample, and cells were incubated at room temperature for 30 minutes. Cells were washed twice with cell staining buffer then resuspended in 200 µL cell intercalation solution comprised of 1X DPBS containing 1.6% paraformaldehyde and 125 nM Cell-ID Intercalator-Ir (Fluidigm). Samples were incubated at 4°C for 12-48 hours then washed sequentially with cell staining buffer and 1X DPBS. To remove clumps of cells, samples were passed through a 40 µm cell strainer (Corning) before a final wash with MilliQ water (Millipore). After the final wash, cells were resuspended in 50 µL MilliQ water for analysis via mass cytometry.

**Mass Cytometry**

Mass cytometry was performed with a Helios mass cytometer (Fluidigm) at the UCLA Jonsson Comprehensive Cancer Center (JCCC) and Center for AIDS Research Flow Cytometry Core Facility. Samples were washed twice with Maxpar cell staining buffer (Fluidigm) and twice with MilliQ water (Millipore). Next, samples were resuspended in 10% EQ Four Element Calibration Beads (Fluidigm) containing natural abundance cerium ($^{140}$Ce/$^{142}$Ce), europium ($^{151}$Eu/$^{153}$Eu), holmium ($^{165}$Ho), and lutetium ($^{175}$Lu/$^{176}$Lu). Samples were run at an event rate of 300–500 events/second, and data was normalized using bead-based normalization in the CyTOF software.

**Mass Cytometry Data Analysis**

Analysis of mass cytometry data was performed in FlowJo V10 (FlowJo LLC). Single, live $CD45^+$ immune cells from each sample in the discovery and validation experiments were manually gated. Each sample was given a unique Sample ID for later deconvolution, then all $CD45^+$ cells from samples in the discovery experiment were concatenated into a single .fcs file. The native FlowJo t-distributed stochastic neighbor embedding (t-SNE) function was used to perform dimensional reduction on the single file containing immune cells from all samples in discovery experiment, using all markers besides CD45 and the following settings: Iterations, 3000; Perplexity, 30; Eta (learning rate), 200. Heat maps of marker expression on the t-SNE plot were generated using the FlowJo Color Map Axis function. Unsupervised k-nearest neighbors (KNN) clustering was performed using the X-Shift algorithm plugin for FlowJo (Samusik et al., 2016). Plugins for FlowJo are found on the FlowJo Exchange and utilize R (R Core Team). X-Shift clustering was performed using all markers besides CD45 and the following settings: number nearest neighbors (K), 54; distance metric, angular; subsampling limit, 42,790; Run ID, auto. Cluster phenotypes were calculated by taking the geometric mean of marker expression by cells in each cluster then transforming using arsinh(x). Immune cell clusters were manually classified as T cells, B cells, NK cells, myeloid cells, and unknown based on expression of the known lineage markers CD3e, CD4, CD8, B220, CD19, CD335, F4/80, CD11b, CD11c, and Ly6G. Phenotypes of immune cell clusters in the discovery experiment were used to manually gate for matched immune cells in the validation experiment. Cluster abundance was calculated by determining the number of cells from each cluster in each sample using the unique Sample ID, then dividing by the total number of $CD45^+$ immune cells in each sample. Age-related immune cell cluster enrichment was analyzed by calculating Z-scores for cluster abundance, using the mean and standard deviation of all replicates and ages for a given cluster. N x N plots were made in FlowJo using the Layout Editor function. Heat maps of cluster phenotype and enrichment were generated in Prism V7.

**Principal Component Analysis**

Principal component analysis (PCA) was performed using immune cell cluster abundance data from the discovery experiment. PCA was performed in R (version 3.6.2) using the singular value decomposition method on scaled matrix data.

**Statistical Analysis**

All statistical tests were performed in Prism V7 (GraphPad). Correlations with age were calculated using the non-parametric Spearman correlation coefficient ($\rho$) with a two-tailed test for significance ($P_\rho$). Comparisons between ages were made using the non-parametric Kruskal-Wallis H test followed by Dunn's multiple comparisons test against the 3-month-old samples, using adjusted p-values to account for multiple comparisons. Kruskal-Wallis and two-way ANOVA were used to evaluate differences in immune lineage markers between immune cell clusters. Fisher's exact test was used to compare the correlation with age ($\rho > 0$ or $\rho < 0$) for identified immune cell clusters between the discovery and validation experiments. Mann Whitney U test was used to compare immune cell cluster frequencies at 3- and 16-months-old for each tissue. Number of replicates (n) and type of replicate are listed in the figure legends. Error bars represent SD. *$p < 0.05$, **$p < 0.01$, ***$p < 0.001$, ****$p < 0.0001$. n.s., not significant, $p \geq 0.05$.
